# Neuronal detection of social actions directs collective escape behavior

**DOI:** 10.1101/2025.09.24.678087

**Authors:** Jo-Hsien Yu, Jimjohn Milan, Geoff T Meyerhof, Julia L Napoli, Matthew Lovett-Barron

## Abstract

Animals in groups obtain information from their social partners to engage in collective behavior^1–4^. Social information transmission has been observed amongst individuals in fish schools^5–8^, bird flocks^9,10^, and human groups^11,12^, but the neural mechanisms for detecting socially transmitted information are poorly understood^4,13–15^. By studying the schooling glassfish *Danionella cerebrum*^16–18^, here we demonstrate that escape from danger is enhanced by visual perception of other escaping fish. We found that neural populations in the midbrain optic tectum^19–21^ and dorsal thalamus^22,23^ are activated by the rapid escape of social partners. These neurons are also driven by the sudden disappearance of virtual social partners, yet unaffected by disappearing stimuli without social relevance. Virtual fish schools that escape or disappear were sufficient to cause observers to escape, even in the absence of direct threats. These results demonstrate that rapid “social-off” detection in visual circuits enables the detection of socially transmitted threat information, which may be a particularly effective strategy for animals capable of rapid movement but limited visual range^17,24^. These results show how neural computations in individuals enables rapid information sharing in animal collectives^4,15^.

## Introduction

Living in groups provides animals with a variety of advantages, including protection from danger and predation^1–10^. In fish schools and bird flocks, an approaching predator or other threatening stimulus can drive collective escape behaviors^7,8,25–29^. Individuals in these groups need not directly sense the threat to avoid danger if they act in response to the movements of their informed neighbors – a benefit of collective living referred to as the “many eyes effect”^30,31^. Despite the prevalence of collective movement and predator avoidance in nature^1–4^, little is known about the neural mechanisms that allow individuals to obtain relevant, actionable information from their partners^4,13–15,32^. The neural basis of social recognition has been characterized in a variety of invertebrate and vertebrate species^18,22,33–36^, but it remains unclear how individuals recognize the specific actions of their social partners to engage in adaptive behavior that benefits the individual and the group.

We address this question by studying the brain and behavior of *Danionella cerebrum*, a micro glassfish that engages in collective schooling behavior using the visual perception of social partners^16,18^ and is amenable to large scale *in vivo* neural activity imaging^18,37,38^. This model system therefore provides us with experimental access to the sensory cues and neural circuits underlying social information transmission during collective responses to threats. Here we examine the perceptual and neural basis of collective escape behaviors in adult *D. cerebrum*, and identify a multi-regional neural signature of social action detection that engages escape behavior in observers to produce collective predator avoidance.

## Results

### Animals in a group escape more effectively than individuals

We first characterized the escape behavior of adult *D. cerebrum* in groups of four^18^, by measuring the positions and postures of fish responding to stimuli that trigger escape behavior (see *Methods*, **Supplemental Fig. 1A**). We applied a mechanical startle stimulus^39,40^ to fish swimming in a 300 mm diameter circular arena, with mechanical taps occurring every seven minutes (see *Methods*; **Fig. 1A**). Taps elicited a short-latency escape behavior (**Fig. 1B, Supplemental Fig. 1B-D**); single fish increased their speed by 10.3±6.1 mm/s in the half second after the tap, whereas a randomly chosen fish in a group of four increased by 31.2±5.8 mm/s (N=15 groups for both). Compared to single fish, fish in groups had greater increase in speed (**Fig. 1C,D**) and escaped to a greater distance after tap onset (**Fig. 1E**). These collective escape behaviors were observed in groups of four or eight fish, but not single fish or groups of two (**Supplemental Fig. 1E-G**). Therefore, the presence of social partners facilitates escape behavior to a mechanical startle.

**Fig. 1.**
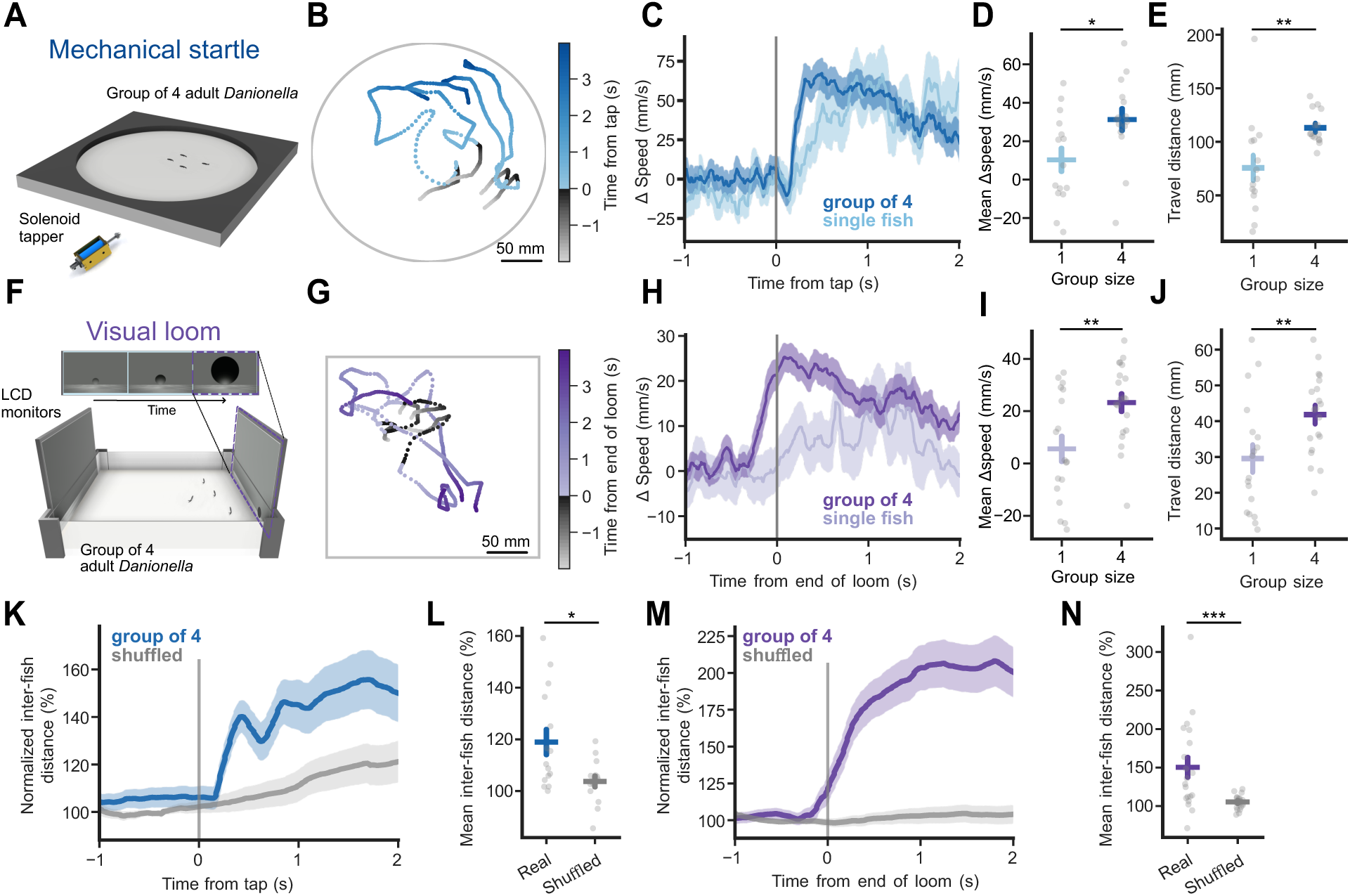
Collective escape behavior of *D. cerebrum* groups. (A) Behavior paradigm for mechanosensation-evoked escape: tap stimuli are delivered by a solenoid actuator hidden underneath the circular arena. (B) Example fish group trajectories over 6 seconds. Each dot is the fish position in a single time frame and is colored by its relative time to tap onset (0 s; gray-to-blue transition). Note rapid change in position after the tap where the consecutive dots are farther apart. (C) Change in speed (mm/s) of fish at tap onset. N=15 groups for single fish and group of 4. (D) Speed increase (mm/s) in 0.5 sec from tap onset. N=15 groups for single fish and group of 4. T-test: t = -2.501, *p* = 0.0185. (10.3±6.1, 31.2±5.8). (E) Travel distance (mm) 1 sec from tap onset N=15 groups for single fish and group of 4. T-test: t = -3.074, *p* = 0.00467. (75.7±11.5, 113.1±4.0). (F) Behavior paradigm for vision-driven escape: looming stimulus is projected by LCD monitors on the wall of the square arena. (G) Example fish group trajectories over 6 seconds. Each dot is the fish position in a single time frame and is colored by its relative time to the end of loom (0 s; gray-to-purple transition). Note the increased change in position started before the stimulus ends, shown as discontinuous dark gray dots. (H) Change in speed (mm/s) of fish at loom offset. N=18 and 20 groups for single fish and group. (I) Speed increase (mm/s) 0.5 sec from end of loom. N=18 and 20 groups for single fish and group. T-test: t = -3.014, p = 0.00469. (5.6±4.9, 23.3±3.4). (J) Travel distance (mm) 1 sec from end of loom. N=18 and 20 groups for single fish and group. T-test: t = -2.574, p = 0.00916. (29.6±3.8, 41.8±2.5). (K) Normalized inter-fish distance (%) at tap delivery. Data from 5 sec before tap onset is used as baseline. Time-shuffled values present in gray. N=15 groups. (L) Changes in inter-fish distance (%) 0.5 sec after tap onset. N=15 groups. Mann-Whitney U = 164.000, *p* = 0.0343. (118.9±4.9, 104.3±2.3). (M) Normalized inter-fish distance (%) at end of loom. Data from 5 sec before end of loom is used as baseline. N=20 groups. (N) Changes in inter-fish distance (%) 0.5 sec after end of loom. N=20 groups. Mann-Whitney U = 338.0, *p* = 1.997 x 10^-4^. (150.3±13.0, 102.4±.9). \**p* <0.05, \*\**p* <0.01, \*\*\**p* <0.001. Data are shown in mean ± SEM.

While mechanical taps are effective in triggering startle behavior and escapes, a more naturalistic threat stimulus is a looming visual object, which mimics an approaching predator. These stimuli drive robust escape and defensive behaviors in aquatic and terrestrial animals^41–45^. We developed a 265 mm x 256 mm square arena to deliver directional looming stimuli to groups of *D. cerebrum*, where two of the four walls were lined with a video screen displaying a smooth-textured floor and dark sky (**Fig. 1F,G**). Looming stimuli were rendered as dark spheres moving towards the center of the arena from one of four randomly selected directions. Stimuli were delivered every seven minutes, with a randomly selected duration between three and six seconds (**Supplemental Fig. 2A**, see *Methods*). Here, we also found that escape behavior was facilitated in groups: single fish escaped by increasing their speed by 5.6±4.9 mm/s in the half second after loom offset, whereas a randomly chosen fish in a group of four increased their speed by 23.3±3.4 mm/s (N=18 and 20 groups for single fish and group of four). Looming stimuli caused a greater increase in speed (**Fig. 1H,I**) and distance travelled (**Fig. 1J**) in fish groups compared to individuals, and these results were also present in larger group sizes (**Supplemental Fig. 2B-D**). Together, these results demonstrate that *D. cerebrum* exhibit a collective escape behavior to visual looming stimuli.

While some dense animal collectives show coherent directional escape “waves” in response to external threats^8,26–28,44^, we observed that fish groups scattered in response to mechanical or visual threats (**Fig. 1K-N**). Groups of four fish increased their mean inter-animal distances upon the onset of each stimulus (tap: 118.9 ± 4.9% of baseline; loom: 150.3 ±12.9% of baseline; **Fig. 1L,N**). Therefore, groups of *D. cerebrum* exhibit a stereotyped collective escape behavior to mechanosensory or visual threats by rapidly scattering, an action commonly referred to as a “flash expansion”^46–48^. This type of collective escape strategy may improve the fitness of group members by preventing predators from following all animals in the group at once^46,49–51^.

### Visual access to social partners enables collective escape

*D. cerebrum* school using their vision of social partners^16,18^, and thus we hypothesized that individuals obtain information from conspecifics using vision to modulate their escape behavior. We tested this hypothesis by quantifying and manipulating visual access during escape behavior. In a separate cohort of fish, we first manipulated the visual cues available to groups during mechanosensory-driven escape (**Fig. 2A**) – removing visible light to prevent fish from interacting with social partners using vision. Single fish exhibited similar escape behavior in the light and darkness to the stimuli (**Supplemental Fig. 3A-C**; N=15 and 13 for light and darkness respectively), indicating that vision had minimal interaction with mechanical startle in individuals. When in groups of four, fish also exhibited similar escape behaviors in both light (Δ88.2±5.1 mm/s) and darkness (Δ80.0±5.4 mm/s) with comparable changes in short-latency speed and travel distance (**Supplemental Fig. 3D-F**). However, groups in darkness showed reduced scattering (**Fig. 2B,C**) and escaped to shorter distances over a longer timescale (3 sec; **Fig. 2D, Supplemental Fig. 3D**). Therefore, vision of social partners facilitates collective scattering in response to a mechanical startle. This is consistent with studies of collective escape in other fish species^7,8,31^.

**Fig. 2.**
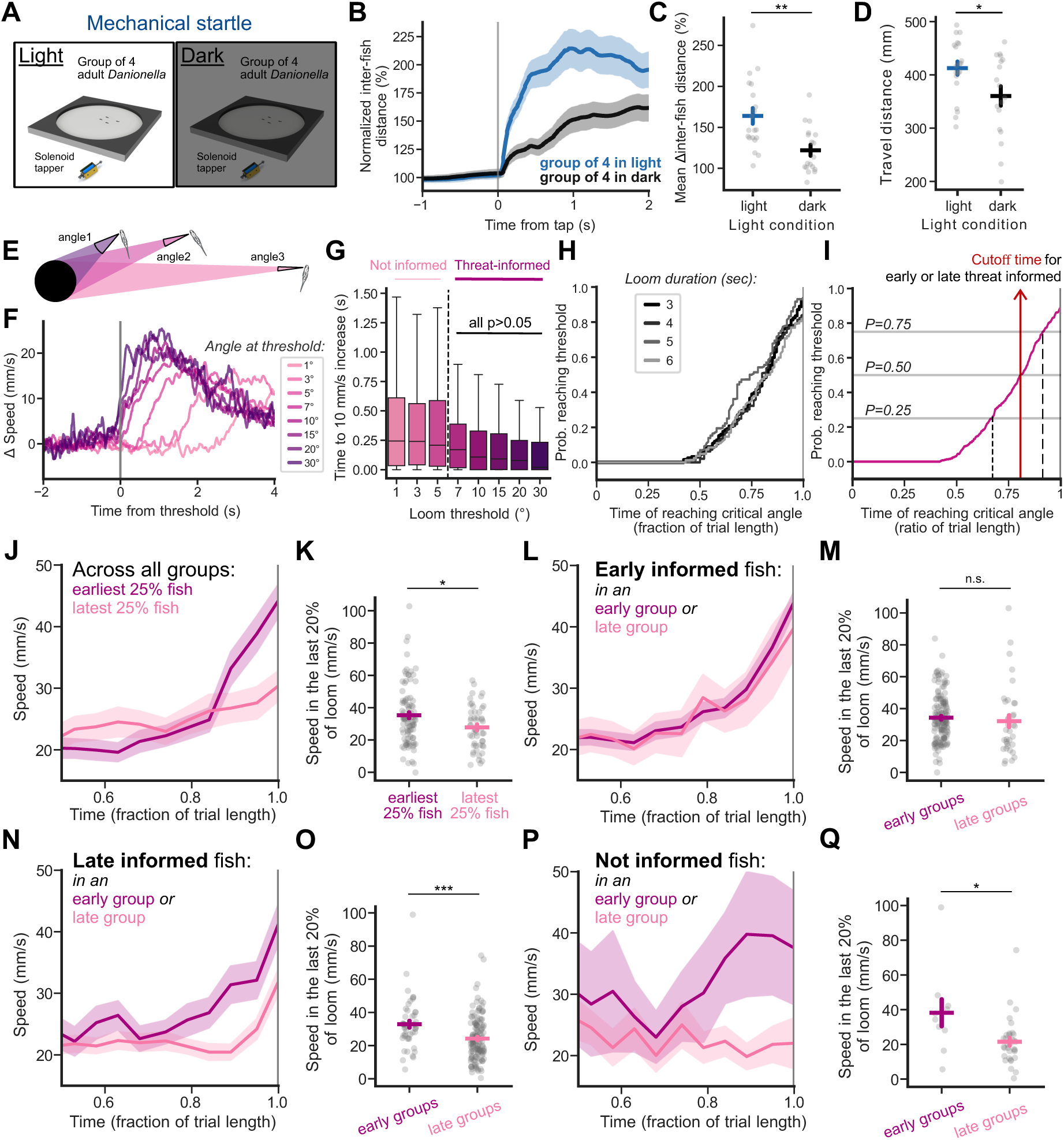
Visual access to escaping social partners drives escape. (A) Behavior paradigm for mechanically-driven escape in light and dark conditions. (B) Normalized inter-fish distance (%) at tap delivery. Data from 5 sec before tap onset is used as baseline. N=20 and 18 groups for light and dark conditions. (C) Changes in inter-fish distance (%) 0.5 sec after tap onset. N=20 and 18 groups for light and dark conditions. T-test t = 3.516, *p* = 0.00120. (164.0±9.6, 122.0±6.7). (D) Travel distance (mm) 3 sec from tap onset. N=20 and 18 groups for light and dark conditions. T-test t = 2.416, *p* = 0.0208. (412.6±12.4, 360.4±18.2). (E) Schematic of eye angles tracing of looming stimulus for multiple fish. Note that the same loom stimulus occupies unequal size in visual field for each fish. (F) Change in speed (mm/s) after crossing visual loom angle thresholds marked with gray line. Speed from 6 sec before the threshold is used as baseline. N=372, 370, 371, 332, 230, 141, 120, 58 for 1°, 3°, 5°, 7°, 10°, 15°, 20°, 30°. (G) Time to 10 mm/s speed increase after crossing different visual loom angle thresholds. N=372, 370, 371, 332, 230, 141, 120, 58 for 1°, 3°, 5°, 7°, 10°, 15°, 20°, 30°. Kruskal-Wallis H = 78.309, *p* = 3.46 x 10^-14^ followed by *post hoc* Dunn’s test with Bonferroni correction. (0.530±0.042, 0.454±0.033, 0.380±0.026, 0.280±0.022, 0.232±0.021, 0.203±0.024, 0.188±0.0026, 0.157±0.032). (H) Cumulative probability of reaching loom angle threshold over time during loom approach for fish separated by the loom duration they experienced. Note the overlap between probability curves. N=126, 96, 72, 82 fish for loom duration of 3, 4, 5, and 6 sec. (I) Cumulative probability of reaching loom angle threshold over time during loom approach across all fish. Time of reaching 0.25, 0.5, and 0.75 are marked with vertical lines. Time to reach 0.5 probability is used to classify fish as early- or late-threat informed in the subsequent analysis. N=376 fish. (J) Speed (mm/s) during loom stimulus of the 25% earliest and latest informed individual across all fish. N=94 and 51 for early and late informed fish. (K) Mean speed (mm/s) of the last 20% of loom stimulus. N=94 and 51 for fish in early and late informed fish. Mann-Whitney U = 2948, *p* = 0.0226. (35.3±1.9, 27.8±1.9). (L) Speed (mm/s) during loom stimulus of “early informed” individuals in early or late informed groups. N=153 and 35 for fish in early and late informed groups. (M) Mean speed (mm/s) of the last 20% of loom stimulus. N=153 and 35 for fish in early and late informed groups. Mann-Whitney U = 3165, *p* = 0.0935. (34.3±1.2, 32.1±3.8). (N) Speed (mm/s) during loom stimulus of “late informed” individuals in early or late informed groups. N=49 and 139 for fish in early and late informed groups. (O) Mean speed (mm/s) of the last 20% of loom stimulus. N=49 and 139 for fish in early and late informed groups. Mann-Whitney U = 4699, *p* = 7.887 x 10^-5^. (32.9±2.1, 24.2±1.1). (P) Speed (mm/s) during loom stimulus of naive individual in early or late informed groups. N=10 and 33 for “not-informed” fish in early and late informed groups. (Q) Mean speed (mm/s) of the last 20% of loom stimulus. N=10 and 33 for naive fish in early and late informed groups. Mann-Whitney U = 255, *p* = 0.0100. (38.9±7.8, 21.5±2.3). \**p* <0.05, \*\**p* <0.01, \*\*\**p* <0.001. Data are shown in mean ± SEM.

While the mechanical threat stimulus is delivered synchronously to all group members, the visual loom stimulus has unequal effects on individual fish, depending on the occupancy of the looming object in their visual field (**Fig. 2E, Supplemental Fig. 2A**); the size of the loom in each individual’s field of view depended on their position in the arena relative to the approach vector (**Supplemental Fig. 4A**). We quantified swimming speed after the loom passed different angular thresholds in each fish’s field of view, ranging from 1 to 30° (**Fig. 2E**, see *Methods*). We observed that speed immediately increases upon passing a threshold of 7°, whereas smaller angles cause delayed escape behavior across different speed of loom approach (**Fig. 2F,G**, **Supplemental Fig. 4B**). Notably, the delay from passing the angular threshold to increasing speed is not significantly different among trials with an angular size threshold of 7° or higher. This critical angle for loom-evoked escape is similar to those reported in adult zebrafish^52^, but smaller than larval zebrafish^42,43,45^.

Within groups of four or more *D. cerebrum*, we examined the angular and temporal distribution of the individuals’ loom angles across the duration of the loom stimulus (**Supplemental Fig. 4C-F**). We calculated the cumulative probability of loom stimulus reaching the 7° critical angle across individual fish and observed a similar temporal pattern (probability reaches 50% at 0.820, 0.801, 0.742, and 0.820 of the normalized loom duration; **Fig. 2H**, **Supplemental Fig. 4G**). We therefor obtained a cutoff time (0.805) for all trials when 50% of fish see the loom at critical angle size or larger (**Fig. 2I**).

We then classified individual fish as “early informed” or “late informed” based on whether that individual’s time to critical loom angle was before or after the cutoff time, respectively (**Fig. 2H, Supplemental Fig. 4H**). While fish increase their speed in advance of the looming stimulus reaching its endpoint, those with earlier sensation of the looming stimulus showed an earlier response and reached faster speeds (**Fig. 2J,K**), regardless of the duration of loom stimulus they experienced (**Supplemental Fig. 4I**).

To determine if individuals benefitted from having early informed social partners, we categorized groups as “early groups” or “late groups” by comparing each group’s average time to reaching critical angle to the same cutoff time. Early-informed fish comprised 72.1% (153/212 fish) of the individuals in early groups, and 16.9% (35/207 fish) of those in late groups. We found that early-informed individuals displayed similar behavior regardless of their group’s status (**Fig. 2L,M, Supplemental Fig. 4J**), suggesting their escape behavior is driven primarily by direct observation of the looming stimulus. In contrast, late-informed individuals increased their speed earlier when they were embedded in early groups, in contrast to those in late groups (**Fig. 2N,O, Supplemental Fig. 4K**). Additionally, fish characterized as “not informed” – those that do not reach the 7° loom threshold – only increased their speed if they were in an early group (**Fig. 2P,Q, Supplemental Fig. 4L**). These results indicate that individuals with limited direct perception of a looming threat stimulus can nonetheless execute a timely escape when their social partners are informed of the threat.

### Fish escape from escaping virtual conspecifics

Our results suggest that fish can escape from threats by responding to their visual perception of escape behaviors in conspecifics. Testing this hypothesis directly requires experimental conditions where single *D. cerebrum* observe conspecifics escaping, but do not sense a direct threat.

To present fish with experimenter-controlled social partners, we developed a visual virtual reality environment populated by 3D fish models whose appearance and movement closely matches those of adult male *D. cerebrum* (**Fig. 3A, Supplemental Fig. 5A,B**, see *Methods*). We found that individual adult fish readily engage with these virtual partners, swimming at an average distance of 71.1±13.3 mm from the monitor, compared to 134.4±17.7 mm at baseline (**Fig. 3B,C**).

**Fig. 3.**
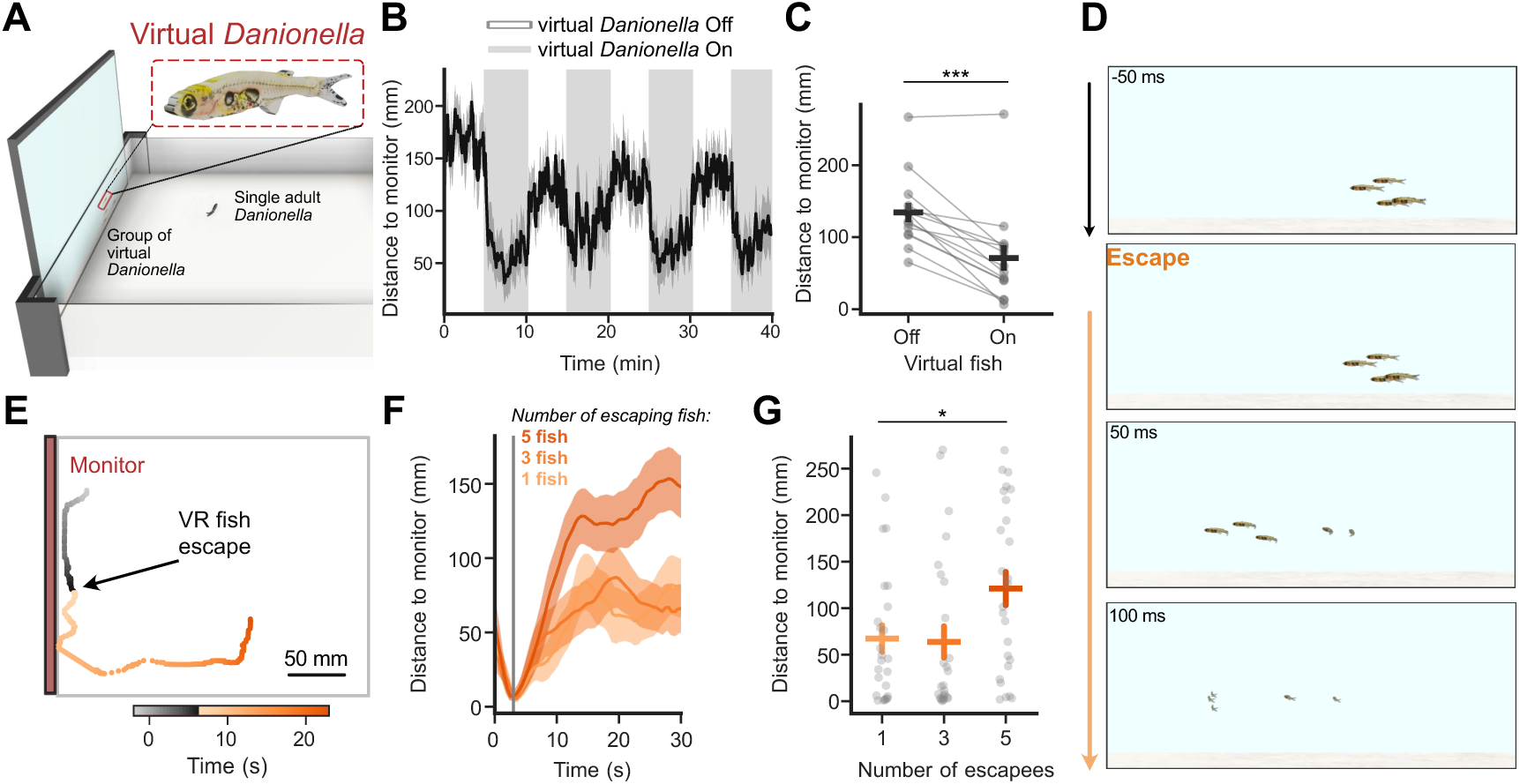
*D. cerebrum* respond to virtual conspecific escape. (A) Behavior paradigm for virtual conspecifics-evoked escape: school of photorealistic *D. cerebrum* is displayed on the LCD monitor on the side of the square arena where a single *D. cerebrum* freely swim in. (B) Distance from the monitor (mm) of the single fish over 40 min. Virtual fish group was rendered during the 10-min blocks in shade. N=13 fish. (C) Mean distance from the monitor (mm) of the single fish. N=13 fish. Wilcoxon signed-rank test statistics = 1.0, *p* = 2.441 x 10^-4^. (134.38±17.679, 71.12±13.333). (D) Example frames of virtual *D. cerebrum* group on the monitor in span of 150 msec at a proximity-triggered escape. Note the scattering of virtual school away from the screen during the escape. (E) Example fish trajectories over 30 seconds. Positions are colored with their relative time to onset of virtual conspecific escape. Note the gray portion of position moving parallel to the monitor before the escape event is triggered. (F) Distance from monitor (mm) of the single fish. Virtual escape onset is marked with a gray line. N=26 fish for trials of 1-, 3-, and 5-fish escaping in a virtual school of five. (G) Distance from monitor (mm) of the single fish 10 sec after the virtual escape onset. N=26 fish for trials of 1-, 3-, and 5-fish escaping in the virtual school of five. Kruskal-Wallis H = 7.730, *p* = 0.0209 followed by *post hoc* Dunn’s test with Bonferroni correction. (67.4±14.3, 63.8±16.7, 121.1±17.9). \**p* <0.05, \*\**p* <0.01, \*\*\**p* <0.001. Data are shown in mean ± SEM.

We developed a closed-loop paradigm where virtual fish would execute an escape action when real *D. cerebrum* came within 16 mm of the virtual groups (> 60 s inter-stimulus interval; **Figs. 3D, Supplemental Fig. 5C**). Because *D. cerebrum* scatter during collective escape (see **Fig. 1K-N**), we reasoned that real fish would commonly escape away from the screen, rather than into it. We observed that individual fish executed rapid avoidance behaviors to these escaping stimuli (**Fig. 3E)**, which was scaled to the number of escaping virtual partners (**Fig. 3F,G**).

Therefore, we conclude that collective escape in *D. cerebrum* is mediated by the visual perception of social partners, where observation of an escaping conspecific promotes one’s own escape. For individual fish to achieve this, their visual systems must be capable of detecting the escape actions of their social partners.

### Neuronal encoding of observed social actions

Neurons in the visual systems of *D. cerebrum* and zebrafish are responsive to stimuli whose size, shape, and motion patterns resemble conspecifics^18,22,53^. These objects robustly activate populations of neurons in the optic tectum (superior colliculus in mammals) – a retinal-recipient midbrain structure that mediates innate visually-driven orienting and movement behaviors in vertebrates ranging from fish to birds to primates^19–21,44,54,55^.

To examine how neural populations in the *D. cerebrum* visual system respond to the escape behaviors of conspecifics, we produced virtual social motion stimuli with routine swimming movements versus those with sudden escape behavior (see *Methods*). With a *D. cerebrum* transgenic line expressing a nuclear-localized calcium indicator in almost all neurons^37,56^, we performed large-scale two-photon calcium imaging from tethered fish within a panoramic visual display environment^18^ (**Fig. 4A**). We used simplified biological motion stimuli^18,22,57^, with fish-sized spheres whose motion in virtual space is derived from recordings of real *D. cerebrum* groups (**Supplemental Fig. 6A**). We presented fish with “social” dots moving in a typical swimming pattern or escape behavior, or objects moving without biological motion – a sphere with linear motion or a looming approach (**Fig. 4B**, see *Methods*). We presented each stimulus for 6 s, in a pseudorandom order with a 15±1 s inter-stimulus interval, and recorded neural activity from 12 fish, which observed each stimulus 7-10 times.

**Fig. 4.**
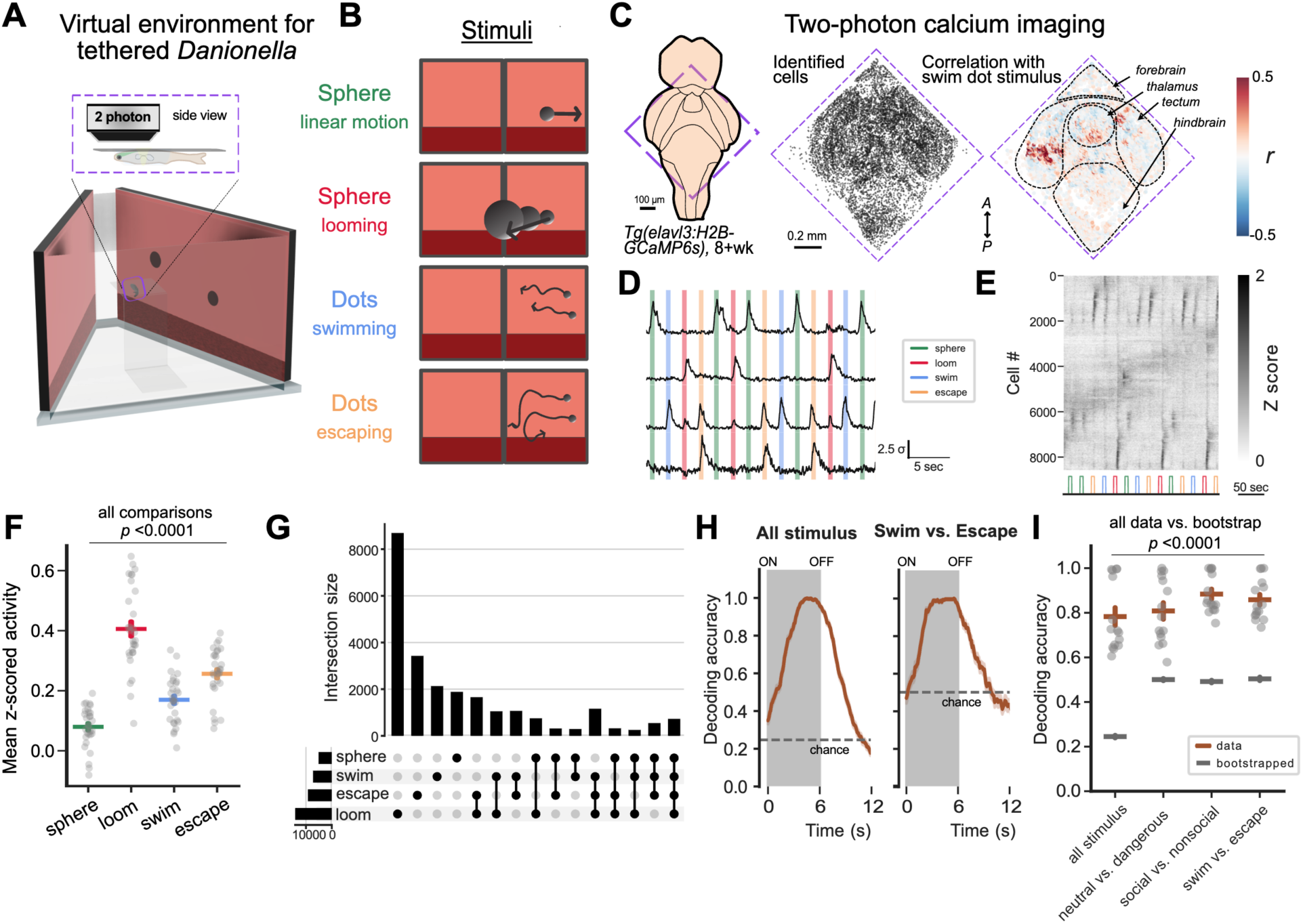
Neuronal encoding of conspecific actions. (A) Schematic of *in vivo* calcium imaging in head-tethered *D. cerebrum* with panoramic visual display of virtual environment. Inset plot shows a more detailed view of fish suspended under coverslip secured in the middle of the arena (see *Methods*). Note that VR display on the monitors encompasses 270° of azimuth and is flattened for ease of visualization in panel B. (B) Illustration of the four types of visual stimuli (top to bottom): linear moving sphere, looming motion sphere, spheres in swimming and escape movements from freely moving *D. cerebrum*. (C) (Left) Dorsal view of adult *D. cerebrum* brain expressing nuclear-localized GCaMP6s. Dashed red line indicates the field of view covering midbrain optic tectum and surrounding brain regions. Positions of all cells extracted (middle) and colored by the correlation coefficient of their activities to the swimming conspecific stimulus, delivered to the right visual field (right); example fish. (D) Short epoch of stimulus-driven single neurons’ responses in an example fish over 30 sec. Color blocks indicated the display time of each stimulus type. (E) Example raster map from cells recorded in (C) aligned with the displayed visual stimuli. Color corresponds to the stimulus types in (B). (F) Maximal Z-scores of cells responsive to each stimulus type. Each dot represents the mean of all cells in one recording session. N=30 sessions. Kruskal-Wallis H = 75.650, *p* = 2.628 x 10^-16^ followed by *post hoc* linear mixed effects model analysis. (0.0796±0.012, 0.405±0.0254, 0.169±0.0141, ±0.256±0.0155). (G) Intersection plot (UpSet) showing response combinations of neurons in all fish recorded. Neural responses overlap across non-biological and biological stimulus types (sphere with linear motion, looming disk, swim (conspecific motion), escape (conspecific motion)). N=23452 cells. (H) Accuracy of support vector machine models over time. Gray shade marks the time of stimulus presentation. Horizontal dashed line is the chance level, calculated from mean accuracy score of models trained with bootstrapped data. From left to right: model trained to classify all four stimulus types, model trained to classify swim and escape stimulus. N=548 and 274 iterations for model for all stimulus and model for swim vs escape stimulus. (I) Mean accuracy of models during stimulus presentation. Each point represents the mean performance of model trained with cells recorded in one session. N=15 sessions. Gray bars represent the average performance of model trained with bootstrapped data across sessions. All stimulus: Mann-Whitney U = 225, *p* = 3.391 x 10^-6^; neutral vs dangerous: T-test t = 8.210, *p* = 6.176 x 10^-9^;social vs nonsocial: Mann-Whitney U = 225, *p* = 3.378 x 10^-6^; swim vs escape: T-test t = 14.180, *p* = 2.637 x 10^-14^. \**p* <0.05, \*\**p* <0.01, \*\*\**p* <0.001. Data are shown in mean ± SEM.

We recorded from 3322±116 neurons in a 978 μm x 978 μm field of view in each of the 50 planes in a total of 12 fish (1-2 planes per imaging session, 1-4 sessions per fish; **Supplemental Fig. 6B**). After motion correction and cell extraction^58^ (**Supplemental Fig. 6C**, see *Methods*), we organized neurons by their anatomical position to identify from neurons in the midbrain optic tectum, as well as surrounding diencephalic and hindbrain cells (**Fig. 4C, Supplemental Fig. 6C**). Many neurons responded to visual stimuli (**Fig. 4D,E, Supplemental Fig. 6D**, see *Methods*), including cells in portions of the optic tectum corresponding to the retinotopic position of stimuli presented to the contralateral eye (**Supplemental Fig. 6E**). We also observed visually-driven cells in the dorsal thalamus, a region identified as responsive to biological motion in juvenile zebrafish^22,23^. We sorted recorded neurons by their responses to visual stimuli, to identify cells with significant responses to individual or combinations of stimuli (**Fig. 4F,G, Supplemental Fig. 6F**; N=23452 neurons in 12 fish, p>0.05). Many cells were primarily activated by the looming stimulus (**Fig. 4F,G, Supplemental Fig. 6D,F,G**); this is expected, as this stimulus grows to occupy most of the visual space (**Supplemental Fig. 6H**), and is innately salient^44^. While many neurons responded to multiple stimuli (**Fig. 4D,F, Supplemental Fig. 6G**), a decoder trained on the principal components of neural activity was able to distinguish amongst all stimulus types (**Fig. 4H,I, Supplemental Fig. 6I-K**, see *Methods*).

We focused our analysis of the smaller stimuli presented to similar positions in visual space: the two social stimuli – routine swimming and escape – as well as the non-biological motion stimulus (**Fig. 4B**). Of these stimuli, the magnitude of neural responses to social swimming and escape were higher than responses to the non-social sphere stimulus (p=3.01 x10^-7^ and p=4.18 x 10^-9^, respectively), and responses to escaping dots were greater than to swimming dots (p = 9.45x 10^-3^; **Fig. 4F**, **Supplemental Fig. 7A**).

Therefore, stimuli of social relevance to the observer boost the response of visually-driven neurons in the midbrain and thalamus. Similar types of gain enhancement have been identified in the tectum/colliculus of other species in response to innately salient visual stimuli^19–21,54^. We hypothesize the gain increases we observe in the *D. cerebrum* visual system are due to the innate salience of social actions, particularly conspecific escape.

### Neuronal detection of vanishing social partners drives escape

To determine how specific social actions are encoded in neural populations, we compared the timing of neural responses to each social stimulus – objects with routine movement versus those with sudden escapes. This analysis revealed a subpopulation of neurons whose activity increased upon the escape action (**Fig. 5A,B, Supplemental Fig. 7A**). When we examined these neurons’ activity during the routine swimming stimulus, we found these cells also responded at offset of the stimulus at trials’ end (**Fig. 5A,B, Supplemental Fig. 7B**). Both neural responses were aligned with the sudden drop in the occupancy of these stimuli in the observer’s visual field (**Supplemental Fig. 6H**). When comparing the time to peak of cells across the two stimuli, we found cells with clear escape-driven activity and offset-driven activity (**Fig. 5C, Supplemental Fig. 7C**). Notably, this stimulus-offset response was specific to social stimuli that move with biological motion; these cells were not driven by the offset of the non-biological translating sphere (**Fig. 5B-D, Supplemental Fig. 7D,E**).

**Fig. 5.**
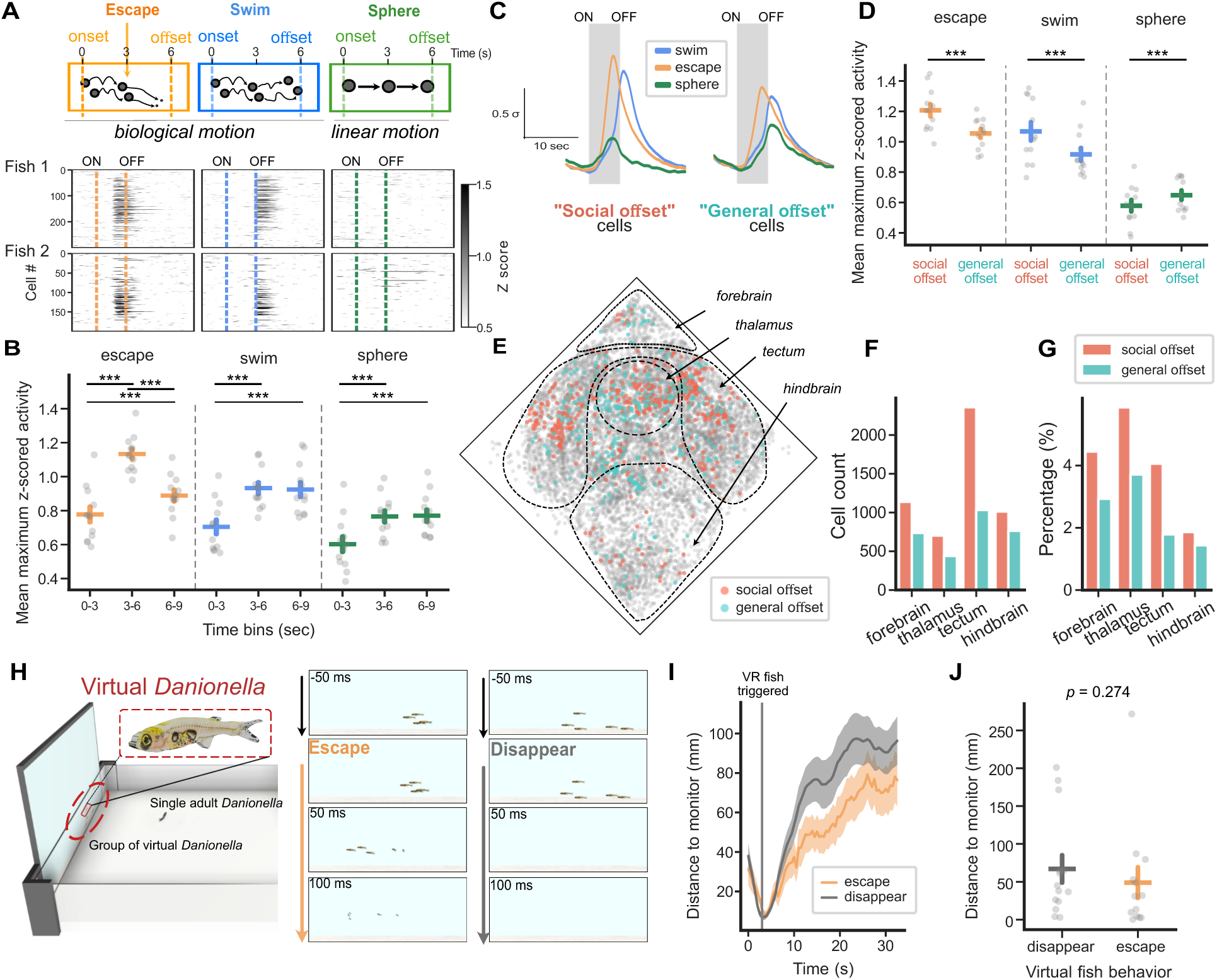
Neuronal detection of social disappearance enables social transmission and collective escape. (A) Top: Schematics of stimulus movements for the social swim, social escape, and non-social sphere during 6 sec of stimulus delivery. Bottom: trial-averaged responses of neurons in two example fish, to each stimulus. Note the timing differences in responses to social swim and escape stimuli, and the lack of response to non-social sphere stimulus. (B) Mean of the maximum z-scored activity of socially responsive cells per fish, grouped by their time-to-peak for each stimulus. Each point is the average of all cells recorded in one fish. N=40375 cells from 12 fish. A linear mixed effects model analysis is used to compare across time within stimulus type (escape: 0.777±0.047, 1.133±0.031, 0.888±0.035; swim: 0.704±0.043, 0.932±0.035, 0.924±0.044; sphere: 0.602±0.046, 0.765±0.035, 0.770±0.036). (C) Left: mean z-scored responses to each stimulus type from all “social offset” cells. N=15711 cells. Right: mean z-scored responses to each stimulus type from all “general offset” cells. N=30444 cells. Gray shades mark the stimulus presentation time. (D) Mean maximum z-scored activity of “social offset” and “general offset” cells to each stimulus type. Each point is the average of all cells recorded in one fish. N=15711 and 30444 cells. Linear mixed effects model analysis to compare response to the same stimulus across cell types and across stimuli within cell type (escape: 1.2080.041, 1.055±0.029; swim: 1.068±0.060, 0.917±0.043; sphere: 0.579±0.038, 0.648±0.031). (E) Positions of all cells extracted in an example fish. “Social offset” and “general offset” cells are colored. Note the different spatial distribution of the two cell types in optic tectum and thalamus. (F) Total cell number across anatomy regions (1137, 702,2357, and 1012 “social offset” cells in forebrain, thalamus, tectum, and hindbrain; 736, 439, 1032, 763 “general offset” cells in forebrain, thalamus, tectum, and hindbrain). (G) Percentage of cells in each anatomical region (4.4%, 5.8%, 4.0%, and 1.8% “social offset” cells in forebrain, thalamus, tectum, and hindbrain; 2.9%, 3.7%, 1.7%, and 1.4% “general offset” cells in forebrain, thalamus, tectum, and hindbrain). (H) Left: behavioral paradigm of virtual escape assay. Right: example frames of virtual *Danionella* group on the monitor in span of 150 ms at a proximity-triggered escape or disappearance event. (I) Mean distance from monitor (mm) of the single fish at virtual escape onset. Gray line shows the escape or disappearance onset. N=13 and 14 fish for virtual escape and disappearance. (J) Mean distance from monitor (mm) of the single fish 10 sec after virtual escape onset. N=13 and 14 fish for virtual escape and disappearance. Mann-Whitney U = 114, *p* = 0.27. \**p* <0.05, \*\**p* <0.01, \*\*\**p* <0.001. Data are shown in mean ± SEM.

We examined the spatial location of “social offset” neurons, responsive to both swim offset and escape but not the sphere offset (**Fig. 5E**, 15711 neurons. N=12 fish). We found 45.2% (2357/5208 total) of these cells in the tectum and 13.4% (702/5208 total) in the dorsal thalamus (**Fig. 5F,G)**. This data suggests that tectal and thalamic neurons tracking the biological motion of social partners are selectively responsive to the disappearance of those partners. As adult *D. cerebrum* are capable of high-speed movements (see **Fig. 1** and ref ^39^), a social partners’ sudden disappearance from their position in the visual field is an indication of their rapid escape.

To determine if a fish’s visual perception of conspecific disappearance impacts their escape behavior, we engaged individual freely-moving *D. cerebrum* with virtual fish models (**Fig. 3A-C, Supplemental Fig. 5**) programmed to suddenly escape or vanish when in close proximity to the real fish (**Fig. 5H**). We compared their behavioral responses to virtual conspecifics executing a kinematically realistic high-speed escape maneuver versus sudden disappearance (**Fig. 5I**, see *Methods*), and observed no difference in escape behavior (**Fig. 5J**).

Therefore, we conclude that individual *D. cerebrum* respond to the escape behavior of their social partners through the visual detection of their sudden disappearance in tectal and thalamic populations. This is effective means of social information transmission that promotes adaptive collective escape behavior at the level of the group.

## Discussion

By analyzing the perceptual and neural basis of escape behavior in *Danionella cerebrum* groups, we have found that the information transmission underlying this collective behavior is mediated by the visual detection of disappearing conspecifics. From our neural recordings, we suggest this is accomplished by populations of visually-responsive neurons in the optic tectum and thalamus, which are activated by the rapid disappearance of social objects with biological motion. This neural function – a “social-off” response – is an effective means for visual animals to detect the escape actions of one’s social partners, and thus execute an escape themselves. Such access to the sensory experiences of social partners (for instance, their detection of a predator) is a major benefit of living in groups, and exemplifies the “many eyes” effect^2,4,30,31^. Our work provides evidence of how neural computation at the level of individual animals can produce emergent collective behaviors at the level of the group^4,14,15^.

Large-scale neural recordings demonstrated the role of the optic tectum in perception of escaping social partners. The tectum/colliculus is known to mediate salience-based attention and neuronal gain across vertebrates^19–21,44,54,55^. While most studies have investigated responses to salient stimuli without social features, socially-relevant visual stimuli are known to activate tectal neurons in fish^18,22^, mice^59^, and primates^55,60,61^. Therefore, the rapid visual detection of social actions with innate salience may be mediated by these conserved circuits across species. Whether the specific “social-off” response emerges from the ON/OFF organization of presynaptic retinal ganglion cells^62^, or is generated by local circuits in the midbrain^19,20^, remains to be determined. We also found neurons for social action detection in the dorsal thalamus, which has been associated with biological motion perception and social behavior in zebrafish^22,23^. Interconnected tectal-thalamic circuits^20,22^ may play an important role in detecting the species-typical movements of social partners to guide collective behavior.

Visual attention to social actions may be particularly important for the adaptive behavior of schooling fish. Aquatic animals possess a more restricted visual range compared to terrestrial animals^24^, and *D. cerebrum* are found in turbid waters with reduced visibility^17^. Therefore, individual animals may be particularly attuned to the actions of their neighbors, whose movements can inform observers of stimuli beyond their individual sensory range. Furthermore, as *D. cerebrum* are capable of high-speed escape movements, an escaping social partner would rapidly leave the short reach of an individual’s field of view. The scattering behavior we observed here may be a consequence of these features, as the socially-transmitted signal for danger (disappearing conspecifics) may not provide the observer with sufficient information about the direction of the threat – only its imminence. For species that show collective “escape waves”^8,26–28,44^, social transmission of threat information may also include information about threat direction. Future studies can examine how group composition, ecology, and neurobiological mechanisms influence the diverse predatory avoidance strategies of animal collectives.

Our results are an instructive example of how the sensory ecology of an organism impacts the behavioral significance of its neural computations and perceptual abilities^63^. *Danionella* will be a useful model clade to investigate this concept^38,64^, as different species occupy distinct ecological niches, such as clear versus turbid waters. Further insights into these natural behaviors will be gleaned by advancing experimental techniques for neural circuit dissection in *Danionella* and other emerging model systems capable of collective behavior^15,65,66^.

## Author Contributions

J-HY, GTM, and ML-B designed experiments; J-HY, JM, and GTM conducted behavioral experiments; J-HY conducted imaging experiments; J-HY, JM, GTM, JLN, and ML-B contributed software and analyzed data; J-HY and ML-B wrote the paper, with assistance from all authors.

## Acknowledgements

We thank Johnatan Aljadeff, Byungkook Lim, Lisanne Schulze, David Zada, and members of the Lovett-Barron Lab for feedback on the manuscript. We thank Princess Tarabishi for pilot experiments and technical assistance, and Mykola Kadobiasnskyi and Benjamin Judkewitz for assistance with anatomical identification. We acknowledge funding from the UC San Diego J. Yang Scholarship (J-HY), the Taiwanese Government Scholarship to Study Abroad Award (J-HY), NIH T32GM133351 (JLN), the Searle Scholars Award (ML-B), Sloan Research Fellowship (ML-B), Packard Foundation Fellowship (ML-B), Pew Biomedical Scholar Award (ML-B), Klingenstein-Simons Fellowship in Neuroscience (ML-B), McKnight Scholars Award (ML-B), and the NIH New Innovator Award DP2EY036251 (ML-B).

---

**Supplemental Fig. 1.**
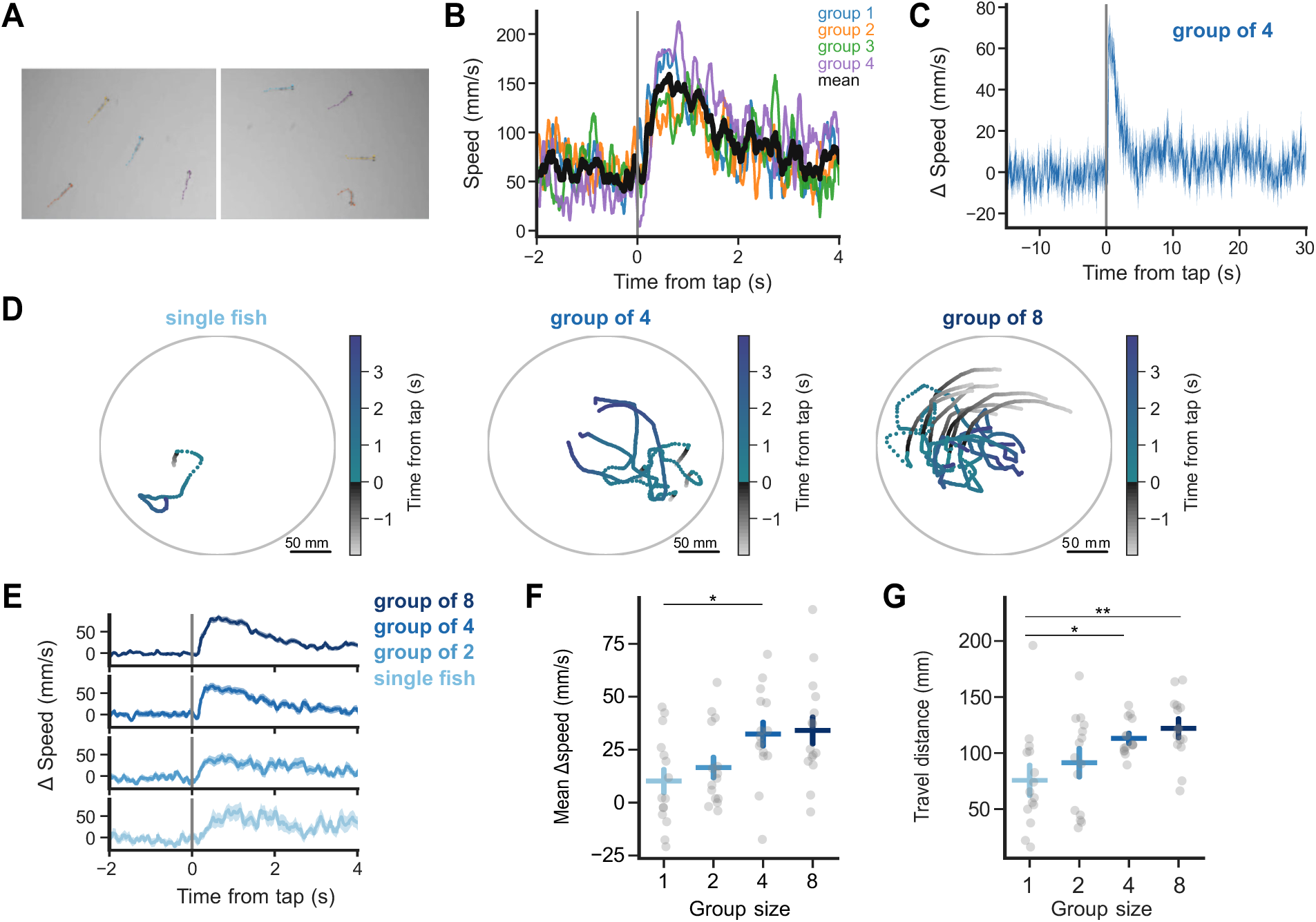
Mechanosensory startle behavior in *D. cerebrum* groups. (A) Example frames with SLEAP inference overlaid for multi-animal tracking. (B) Example speed at tap onset. Data from 4 groups of four and the average are shown in colored and black traces respectively. (C) Change in speed (mm/s) of fish at tap onset. N=15 groups. Noted the punctate escape response to mechanical stimulus in this longer time series. (D) Example fish group trajectories over 6 seconds. Each dot is the fish position in a single time frame and is colored by its relative time to tap onset. From left to right: single fish, group of 4, and group of 8. (E) Change in speed (mm/s) of fish at tap onset. N=15 groups for single fish, group of 2, group of 4, and group of 8. (F) Speed increase (mm/s) in 0.5 sec from tap onset. N=15 groups for single fish, group of 2, group of 4, and group of 8. Kruskal-Wallis H = 11.477, *p* = 0.00940 followed by *post hoc* Dunn’s test with Bonferroni correction. (10.160±5.503, 16.583±4.889, 32.342±5.653, 34.076±6.401) (G) Travel distance (mm) 1 sec from tap onset. N=15 groups for single fish, group of 2, group of 4, and group of 8. One-way ANOVA F = 13.844, *p* = 4.5 x10^-4^ followed by *post hoc* Tuckey’s test with Bonferroni correction. (75.654±11.522, 91.363±10.873, 113.117±3.975, 122.106±4.275) \**p* <0.05, \*\**p* <0.01, \*\*\**p* <0.001. Data are shown in mean ± SEM.

**Supplemental Fig. 2.**
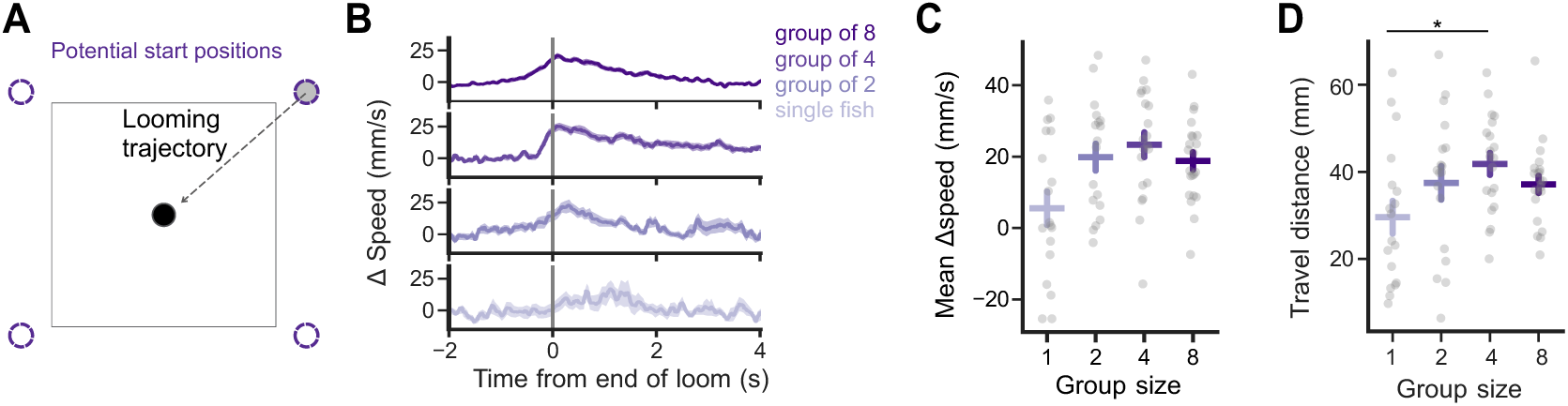
Visual loom-driven escape behavior in *D. cerebrum* groups. (A) Schematics of randomized loom stimulus approach trajectory (B) Change in speed (mm/s) of fish at end of loom. N=18, 18, 20, 22 groups for single fish, group of 2, group of 4, and group of 8. (C) Speed increase (mm/s) in 0.5 sec from end of loom. N=18, 18, 20, 22 groups for single fish, group of 2, group of 4, and group of 8. One-way ANOVA F = 2.792, *p* = 0.0987. (5.555±4.681, 19.857±3.765, 23.354±3.463, 18.783±2.456) (D) Travel distance (mm) 1 sec from end of loom. N=18, 18, 20, 22 groups for single fish, group of 2, group of 4, and group of 8. Kruskal-Wallis H = 8.810, *p* = 0.0319 followed by *post hoc* Dunn’s test with Bonferroni correction. (29.552±3.767, 37.450±3.849, 41.830±2.531, 37.125±1.983) \**p* <0.05, \*\**p* <0.01, \*\*\**p* <0.001. Data are shown in mean ± SEM.

**Supplemental Fig. 3.**
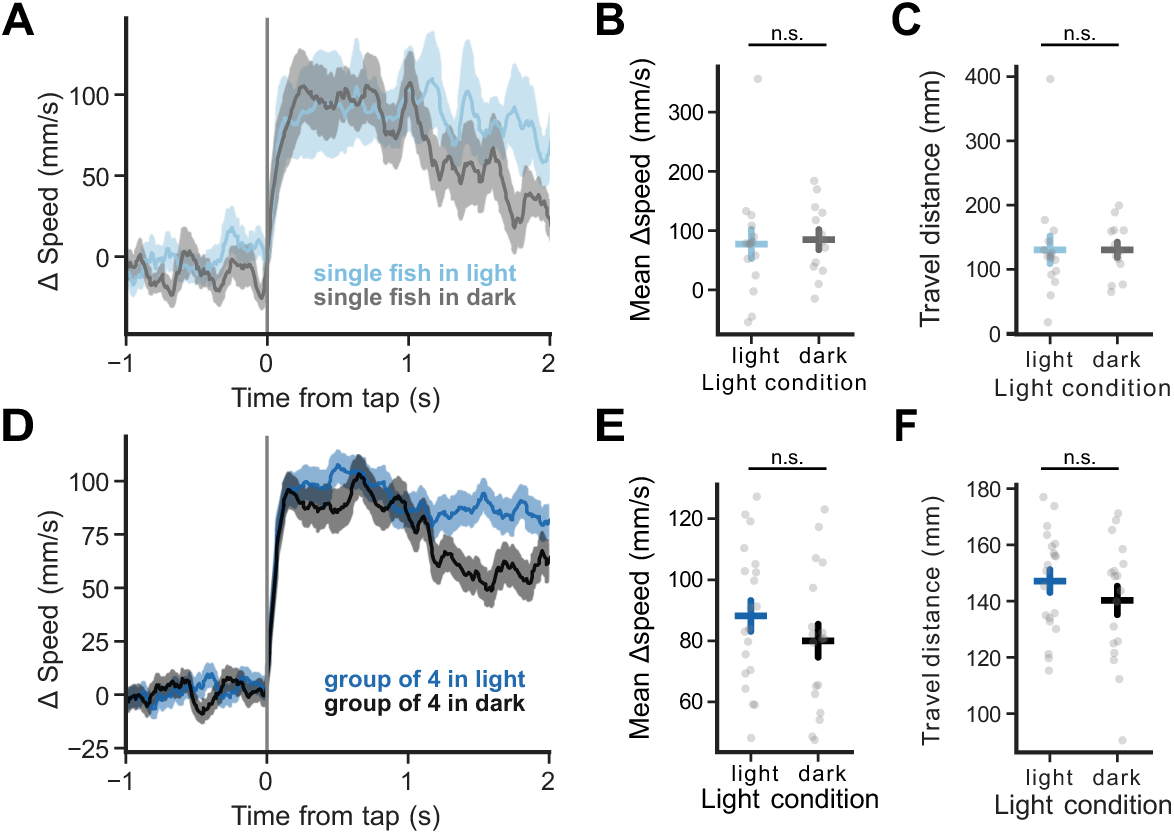
The role of vision in escape to mechanical startle. (A) Change in speed (mm/s) of fish at tap onset. N=15 and 13 single fish in light and dark conditions. (B) Speed increase (mm/s) in 0.5 sec from tap onset. N=15 and 13 single fish in light and dark conditions. Mann-Whitney U = 84.0, *p* = 0.549. (77.209±24.580, 84.558±17.031). (C) Travel distance (mm) 1 sec from tap onset. N=15 and 13 single fish in light and dark conditions. Mann-Whitney U = 83.0, *p* = 0.518. (130.434±21.506, 130.651±11.882). (D) Change in speed (mm/s) of fish at tap onset. N=20 and 18 groups in light and dark conditions. (E) Speed increase (mm/s) in 0.5 sec from tap onset. N=20 and 18 groups in light and dark conditions. T-test t = 1.097, *p* = 0.279. (88.155±5.060, 80.031±5.414). (F) Travel distance (mm) 1 sec from tap onset. N=20 and 18 groups in light and dark conditions. T-test t = 1.072, *p* = 0.290. (147.128±4.067, 140.223±5.071). \**p* <0.05, \*\**p* <0.01, \*\*\**p* <0.001. Data are shown in mean ± SEM.

**Supplemental Fig. 4.**
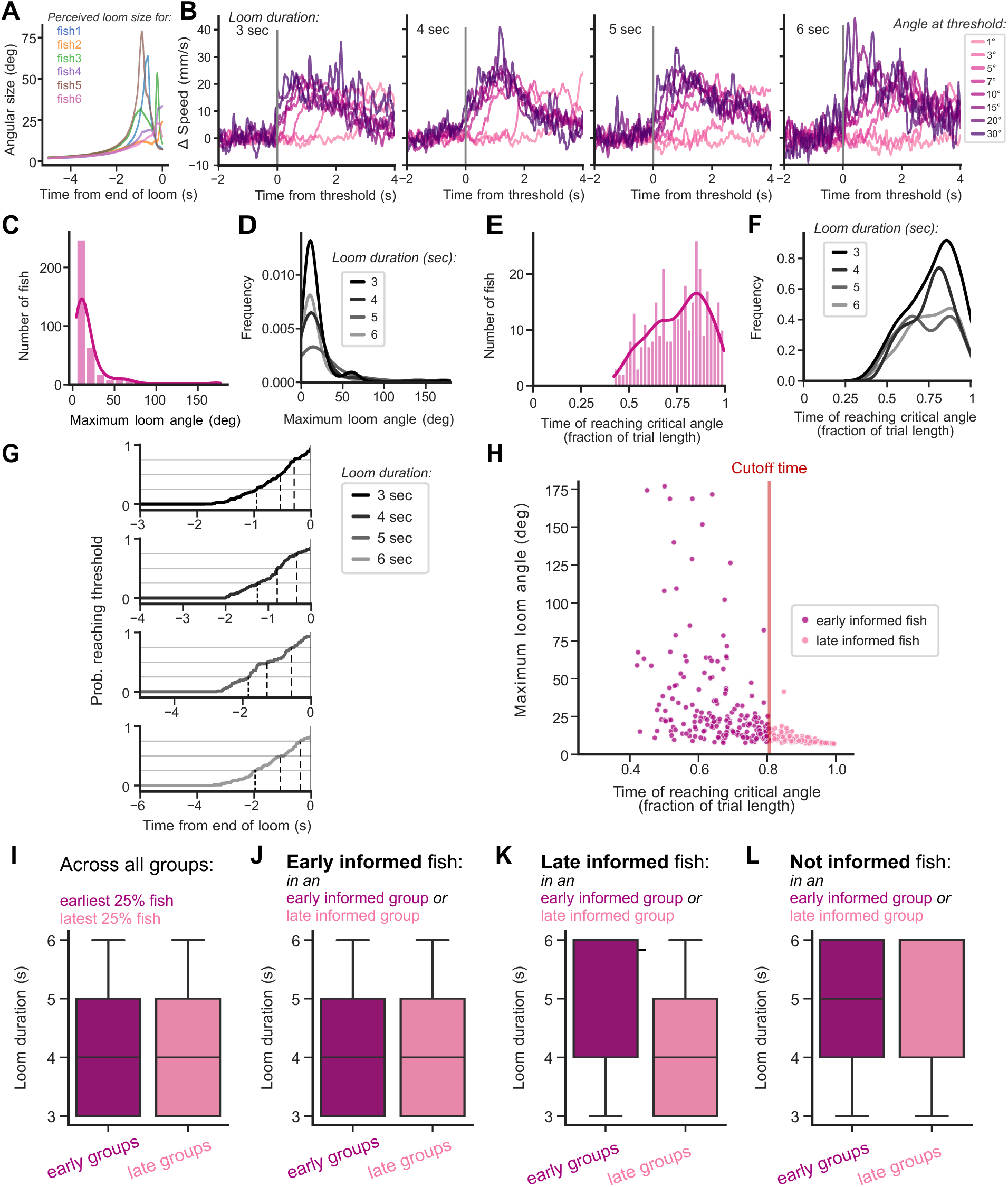
Visual basis of collective escape movements to looming stimuli. (A) Example eye angle traces of loom stimulus for fish in a group over 5 sec. Note that threat proximity varies across fish in the group. (B) Change in speed (mm/s) after crossing visual loom angle thresholds marked with gray line. Speed from 6 sec before the threshold is used as baseline. From left to right: trials with loom duration of 3, 4, 5, and 6 sec. (C) Distribution of maximum loom visual angle perceived by individual fish. N=376 fish. (D) Frequency distribution of maximum loom visual angle perceived by individual fish separated by loom durations. N=126, 92, 72, 82 fish for loom duration of 3, 4, 5, and 6 sec. (E) Distribution of time for loom stimulus reaching critical angle for each individual fish (ratio of trial length). N=376 fish. (F) Frequency distribution of time for loom stimulus reaching critical angle for each individual fish separated by loom durations (ratio of trial length). N=126, 92, 72, 82 fish for loom duration of 3, 4, 5, and 6 sec. (G) Cumulative probability of reaching loom angle threshold over time during loom approach. Time for the reach 0.25, 0.5, and 0.75 are marked with vertical dashed lines for each curve. From top to bottom: N=126, 92, 72, 82 fish for loom duration of 3, 4, 5, and 6 sec. (H) Scatterplot of maximum loom angle (deg) and time of reaching critical angles for all individual fish. Critical angle and 0.5 probability are shown in vertical lines N=376 fish. (I) Loom duration (s) for the 25% earliest and latest informed individual across all fish. N=94 and 51 for fish for early and late informed fish. Mann-Whitney U = 2693, *p* = 0.204. (4.404±0.116, 4.157±0.162). (J) Loom duration (sec) for the early informed individual in early or late informed groups. N=153 and 35 for fish in early and late informed groups. Mann-Whitney U = 2873.5, *p* = 0.485. (4.333±0.090, 4.200±0.200). (K) Loom duration (sec) for the late informed individual in early or late informed groups. N=49 and 139 for fish in early and late informed groups. Mann-Whitney U = 4275.5, *p* = 5.755 x 10^-3^. (4.633±0.153, 4.151±0.100). (L) Loom duration (sec) for naive individual in early or late informed groups. N=10 and 33 for naive fish in early and late informed groups. Mann-Whitney U = 193.5, *p* = 0.398. (4.900±0.379, 4.545±0.200). \**p* <0.05, \*\**p* <0.01, \*\*\**p* <0.001. Data are shown in mean ± SEM.

**Supplemental Fig. 5.**
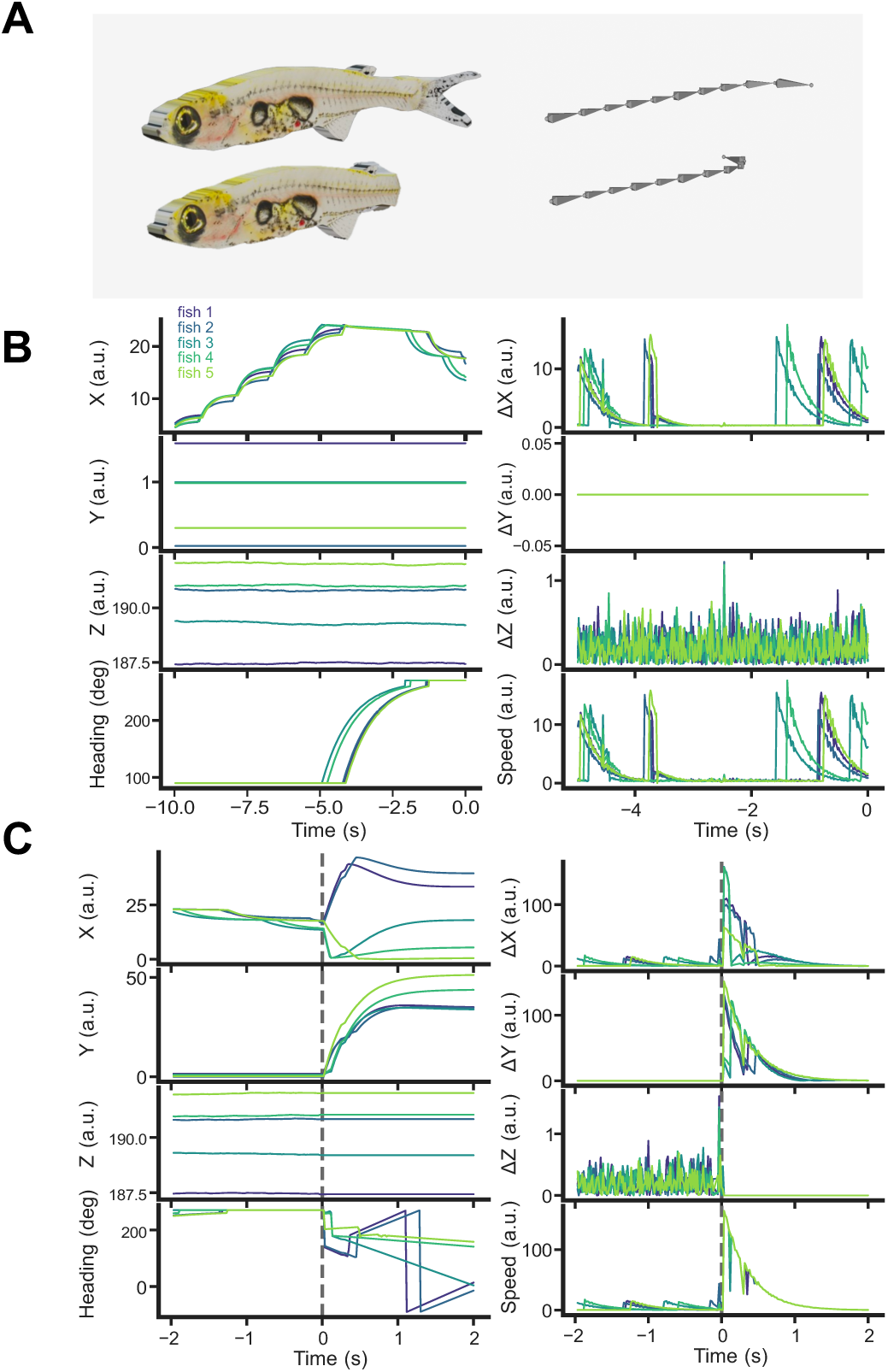
Movements of virtual *D. cerebrum*. (A) *D. cerebrum* model appearance (left) and skeleton (right) used in the virtual reality environment. Note the bending of tail in the model follows the curvature of skeleton in three-dimensional space. (B) Example kinematics of the virtual *D. cerebrum* in the VR environment over 10 sec of normal swimming. Left column from top to bottom: X-, Y-, Z-positions and heading direction. Right column from top to bottom: derivatives of X, Y, Z, and speed. (C) Example kinematics of the virtual *D. cerebrum* in the VR environment over 4 sec at the onset of virtual escape (red dashed line). Left column from top to bottom: X-, Y-, Z-positions and heading direction. Right column from top to bottom: derivatives of X, Y, Z, and speed.

**Supplemental Fig. 6.**
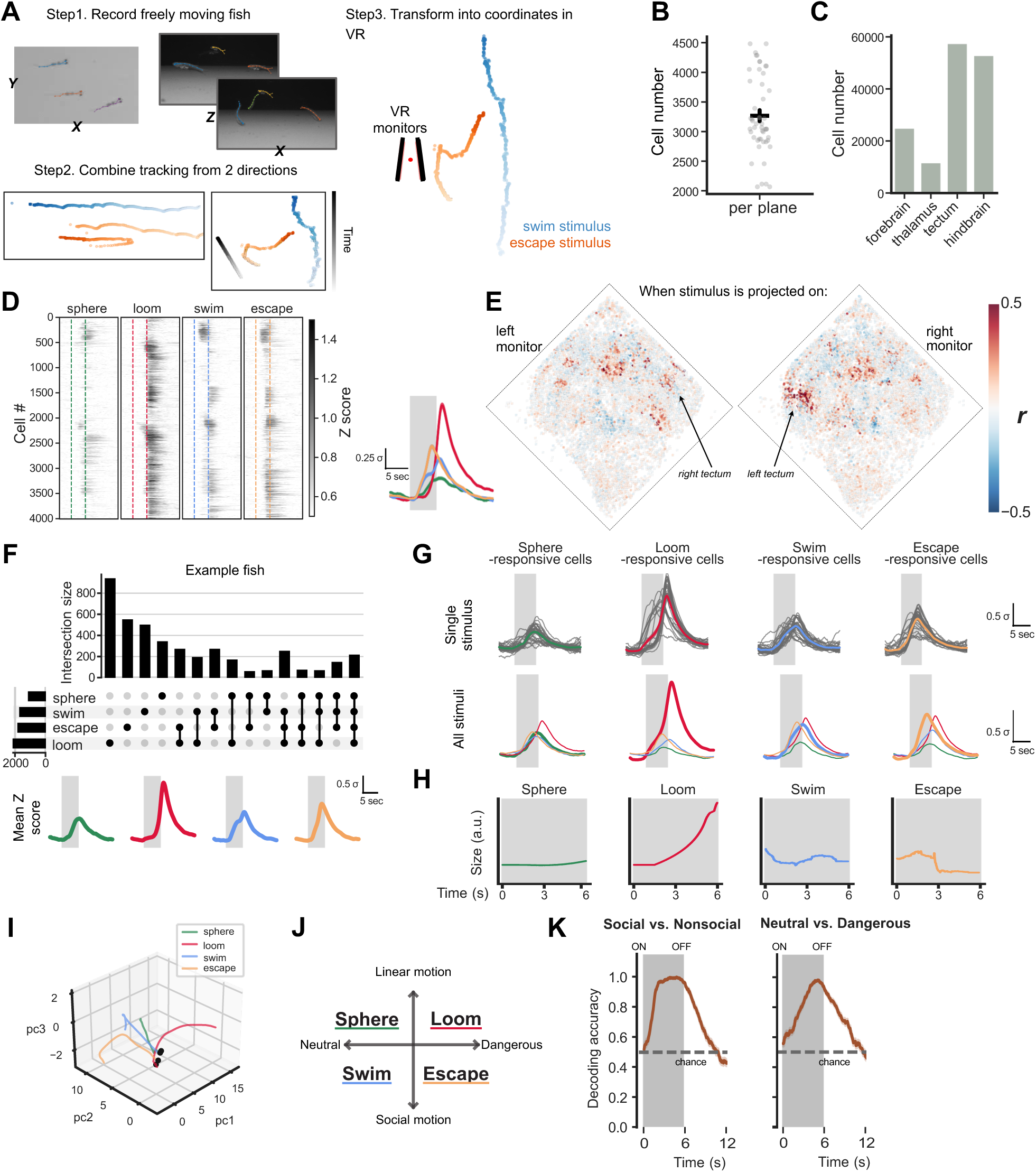
Neural responses to social and non-social visual stimuli. (A) Workflow of making VR social stimuli from real fish movements. (B) Mean cell number recorded per imaging plane. N=50 planes from 12 fish. (3267.1±92.9). (C) Total cell number recorded across anatomical regions. N=25101, 11823, 57649, 53040 cells in forebrain, thalamus, tectum, and hindbrain. (D) Mean z-scored response to each stimulus type from all responsive cells in an example fish. The average neural response is overlaid on the right with gray shade marking stimulus presentation time. N=4006 cells. (E) Positions of all cells extracted in an example fish and colored by the correlation coefficient of their activities to the swimming stimulus during two recording sessions where the same stimulus is on VR monitors of the opposite side. Note the lateralization of highly correlated neuronal clusters on the contralateral side of the visual stimulus. (F) Neural responses overlap across stimulus types in an example fish. N=4006 cells. (G) Mean z-scored response to different stimulus types in all fish recorded. Top: Gray lines show the mean activity of all cells recorded in one session, and their average in colored line. Bottom: thick lines show the mean response to the cell responsive to each stimulus; thin lines show the response of the same cells to other stimulus types. N=13827, 30556, 17871, and 20590 sphere-, loom-, swim-, and escape-responsive cells from 30 sessions in 12 fish. (H) Visual occupancy of different stimulus types. (I) Example low-dimensional trajectory of trial-averaged stimulus responses, beginning at 1 sec pre-stimulus onset (black dot) and progressing for 6 s until stimulus offset. Trajectories are colored by stimulus type and are shown for the first three principal components (PCs). (J) Design of four visual stimuli along two axes: nonsocial or social (linear or biological motion) & neutral or dangerous. (K) Accuracy of support vector machine models over time. Gray shade marks the time of stimulus presentation. Horizontal dashed line is the chance level, calculated from mean accuracy score of models trained with bootstrapped data. From left to right: model trained to classify social vs nonsocial stimulus types, model trained to classify neutral vs dangerous stimulus. N=548 iterations for both models. \**p* <0.05, \*\**p* <0.01, \*\*\**p* <0.001. Data are shown in mean ± SEM.

**Supplemental Fig. 7.**
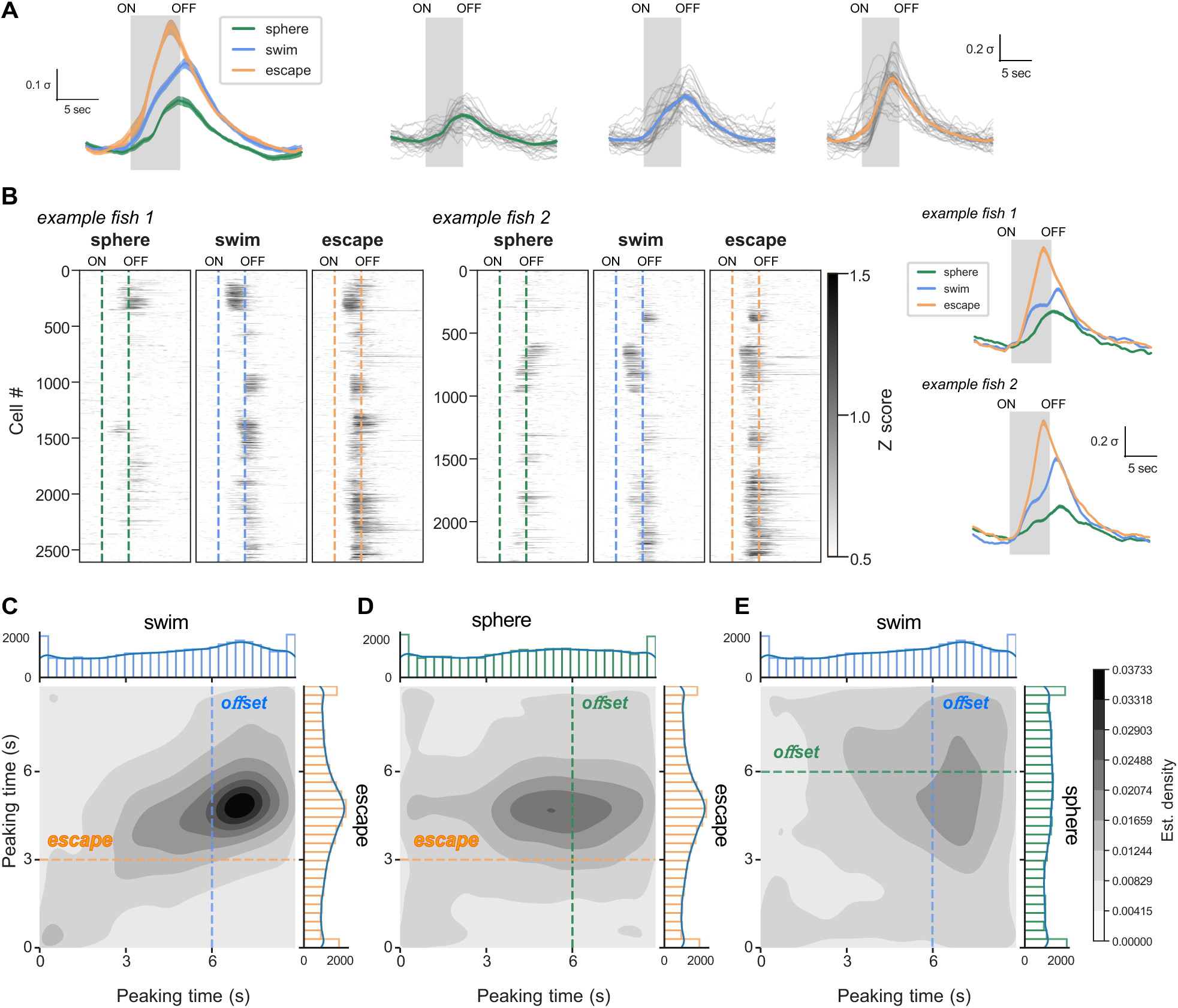
Further characterization of social offset responses. (A) Left: mean z-scored response to different stimulus types in all fish recorded. Right: gray lines show the mean activity of all cells recorded in one session, and their average in the colored line. N=40375 cells from 30 sessions in 12 fish. (B) Mean z-scored response to each stimulus type from all responsive cells in two example fish. The average neural response is overlaid on the right with gray shade marking stimulus presentation time. N=2610 and 2322 cells in the two fish. (C) Density maps (smoothed scatterplot) of peaking time (s) toward swim vs escape stimulus in all cells responsive to small stimuli. Time of escape and swim stimulus offset are indicated with colored lines. N=40375 cells. (D) Density maps (smoothed scatterplot) of peaking time (s) toward sphere vs escape stimulus in all cells responsive to small stimuli. Time of escape and sphere stimulus offset are indicated with colored lines. N=40375 cells. (E) Density maps (smoothed scatterplot) of peaking time (s) toward swim vs sphere stimulus in all cells responsive to small stimuli. Time of swim offset and sphere offset are indicated with colored lines. N=40375 cells. \**p* <0.05, \*\**p* <0.01, \*\*\**p* <0.001. Data are shown in mean ± SEM.

## Methods

### Fish husbandry

*Danionella cerebrum* were raised and maintained in conventional zebrafish housing systems (Aquaneering; system water temperature 29 ± 0.5 °C, pH 7.0, conductivity 650 mS), under 14/10h light-dark cycles, and fed 2-3 times a day. *D. cerebrum* were bred in communal tanks of ∼40 individuals, enriched with (plastic green thing) and ∼5 cm silicone tubes to encourage spawning (citation). Eggs were collected during the first hour of daylight, and 1-2 h after morning and afternoon feeding. Embryos (0 to 5-day post fertilization) were raised in egg-water (0.2 mg/L Instant Ocean, 3 g/l mm CHNaO3, and 0.15% methylene blue dissolved in reverse osmosis purified water) in an incubator at 29 ± 0.5°C.

Larvae (5-14 dpf) were housed in static tanks in the housing systems and cocultured with L-type rotifers (*Brachionus plicatilis*), while water level was raised ∼1 cm per day. Between 14 to 21-day post fertilization, housing tank was connected to the water circulation system and fed with artemia. At 8-10 weeks post fertilization, *D. cerebrum* reach adulthood and become fertile. Wild-type *D. cerebrum* were provided by Dr. Adam Douglass (University of Utah) and Dr. Benjamin Judkewitz (Charité Berlin). The transgenic line *Tg(elavl3:H2B-GCaMP6s)* (ref.^56^) was provided by Dr. Benjamin Judkewitz (Charité Berlin).

### Behavioral experiments setups

All behavioral experiments used 8 weeks or older *D. cerebrum*, and both sexes are included. Behavioral experiments were conducted in a room with a portable electric heater to maintain water temperature between 28-30°C. Data acquisition was automated and timestamped with prebuilt packages and custom workflows in BonsaiRx^67,68^ (bonsai-rx.org). All behavioral arenas were built with custom cut acrylic (black walls and transparent bottom with white styrene to diffuse bottom lit light sources) and encircled with black curtain to control luminance. Light and dark environments are controlled with a bottom-projected white or dark background (AnyBeam Pico Projector, HD301M1-H2).

Behaviors were illuminated with infrared light (CM-IR130-850NM, CMVision Technologies Inc.) from the bottom, and acquired at 121 Hz from a camera mounted above (FLIR Grasshopper, #33-534) with an 8 mm/ F 1.8 lens (Edmund Optics, #15-626) and long-pass filter (Edmund Optics, #12-767).

### SLEAP tracking and Python processing

Social LEAP-Estimates Animal Poses (SLEAP)^69^ was used to train models for tracking the location and posture of *D. cerebrum*. Each video was proofread after inference and identity tracking to create h5 files that contain positional information of all tracking nodes of each animal for the given behavioral session. H5 files and corresponding timestamped csv were processed with custom Python codes for further behavioral analysis (see Code Availability). Models were trained with a 9-node skeleton covering the two eyes and evenly distributed points along the midline of the body.

For each time point, animal centroid was defined as the x/y coordinates of the middle point of the two swim bladders. A nose point was defined as the center point of the two eyes, and the orientation of the animal was defined as the direction of the vector pointing from the centroid to the nose point and smoothed by a 5 -frame running average (∼40ms). Speed and angular speed of the animal was calculated by taking the derivatives of centroid coordinates and the heading direction and smoothed by a 10-frame running average (∼80 ms), respectively. Distance and alignment of each fish to all its neighbors were identified for each time point. All x & y coordinates were then transformed from pixel to mm by taking measurement of arena edge of known physical length in Fiji. Multiple csv files were saved with the timestamps for each video frames and any stimulus presented. Custom Python codes align all arrays to the behavior before further analysis.

### Mechanosensory tap-evoked escape assay

In the tap evoked escape behaviors, *D. cerebrum* were left to habituate for 20 min in a 300 mm circular arena filled with 1 L of system water^18^ in the beginning of the experiment. Each experiment consisted of five 2-minute recordings, where an actuator (model) mounted beneath one corner of the arena was moved by a 300-ms voltage pulse that delivers an intense mechanical wave at the 1-minute mark. There are 7 minutes of intervals between each recording, and groups of 1, 2, 4, and 8 fish were used. For the light-dark experiments of tap escape, groups of 1 and 4 fish from a different cohort were used. Behaviors were run with either a white or black background bottom projection.

Mean of the speed in the 30 sec pre tap onset was used as the baseline for each individual fish to calculate how their speed change at 0.5 sec after stimulus (**Fig. 1D, Supplemental Figs. 1F, 3B, E**).

Travel distance after stimulus was calculated with integration of speed over 1-sec (**Fig. 1E, Supplemental Figs. 1G, 3C,F**) or 3-sec (**Fig. 2D**) window. Inter-fish distance was calculated for each pair of fish within a group. The positions of each fish in the group were shifted by a random number independently to create the shuffled data used to calculate chance inter-fish distance. Change in inter-fish distance was computed by subtracting the mean over 1-sec before tap onset, and the mean at 0.5 sec after the stimulus are used for statistical comparison (**Figs. 1L, 2C**).

### Visual loom-evoked escape assay

In the loom evoked escape behaviors, *D. cerebrum* were left to habituate for 20 min in a square arena (265 x 265mm) where two of the walls were lined with 10.1” LCD monitors (HAMTYSAN, 10.1 inch rendered at 1024x600 pix). Each screen was mapped as a 14x20 cm window into a 3D virtual environment, eye perspective at center point of area, 2 cm above the bottom) using BonVision package^68^. Each experiment consisted of five 2-minute recordings, and at 1-minute mark a dark sphere of 4.3 cm would appear in one of the corners (x=±30, y=±15, z=1.5 cm) and move linearly to center of arena (x=0, y=0, z=1.5 cm) within 3-6 seconds. Both the starting position and moving duration were randomly rendered. There are 5 minutes of intervals between each recording. Groups of 1, 2, 4, and 8 fish were used.

Mean of the speed in the 30 sec pre loom stimulus onset was used as the baseline for each individual fish to calculate how their speed change at 0.5 sec after stimulus (**Fig. 1I, Supplemental Fig. 2C**). Travel distance after stimulus was calculated with integration of speed over 1-sec (**Fig. 1J, Supplemental Fig. 2D**) window. Inter-fish distance was calculated for each pair of fish within a group. The positions of each fish in the group were shifted by a random number independently to create the shuffled data used to calculate chance inter-fish distance. Change in inter-fish distance was computed by subtracting the mean over 1-sec before loom onset, and the mean at 0.5 sec after the stimulus are used for statistical comparison (**Fig. 1N**).

Eye angles tracing of loom was done with two-dimensional ray-casting between the middle point of the two eyes and the loom stimulus for each fish during all time points loom stimulus was present (**Fig. 2E,F, Supplemental Fig. 4A,B**). From the recorded centroid of virtual sphere, we calculated the coordinates of the edges to get their alignment to the fish respectively. The absolute difference between the two angles was the loom angular size.

After identifying the 7° critical angle, we created Boolean arrays to capture when each fish first perceives the loom. These arrays were resampled to align the time of different trials to construct the cumulative probability curve of reaching loom threshold (**Fig. 2H,I**). Time points (in ratio of the trial length, aka loom duration) when the probability curve exceeds 50% was the cutoff time to assign individual and group information status in the subsequent analysis. For individuals, they were labeled as “early informed” if loom reaches critical angle at an earlier time than the cutoff. Otherwise, individuals were labeled as “late informed”, including those never reached critical angles and didn’t have a valid reaching time. Mean of the reaching time in a group was also compared to the same cutoff to classify them as “early group” or “late group”. To compare behavior across trials, average speed in time bins 5% of the original trial length was taken.

### Virtual conspecifics engagement and social escape assay

A 3D virtual reality (VR) environment using Panda3D (https://www.panda3d.org/), and populated environment with five virtual male *D. cerebrum* (**Supplemental Fig. 5A**) modeled in Blender (v. 3.5.1). The VR scene was captured by a virtual camera positioned in VR space such that the virtual *D. cerebrum* appeared on the bottom third of the monitor, and the virtual conspecifics was programmed to appear on one of the monitors, with the other monitor displaying an empty background. Each VR *D. cerebrum* appeared ∼12 mm in length on the monitor and had animated tail movements that coincided with the burst phase of their burst-and-glide kinematics (**Supplemental Fig. 5B**). The virtual school was programmed to swim along the width of the monitor by translating their X position in VR space, with small, random deviations in Z to simulate slight changes in depth. The school always remained in the field of view of the VR camera, and therefore appeared on the monitor, as each fish executed a 180° turn when they reached the boundary of environment.

The environment was rendered on the monitors of the same square arena used in loom evoked escape assay (**Fig. 3A**). To establish whether the sight of our virtual school attracted real fish, we first implemented an open loop escape paradigm (**Fig. 3B,C**). Single *D. cerebrum* was left to habituate for 5 min in a square arena before being shown four times of 5-minute no stimulus followed by 5-minute stimulus-on with five swimming virtual *D. cerebrum*. At the end of each stimulus-on period, all virtual fish were assigned a random escape heading spanning 150° towards the orientation of the virtual camera so they rapidly swam away from the virtual camera into the horizon of the VR environment (**Fig. 3D**).

For the social escape assays, we implemented a closed-loop paradigm wherein the proximity of the real fish to the virtual school triggered an escape. For this, we again allowed a single *D. cerebrum* to habituate the environment for 5 minutes. After 1 minute of the 5-fish virtual school being displayed on the monitor, the test fish’s engagement with the virtual school (test fish distance to center of virtual school < 16 mm) triggered either an escape or disappearance of VR fish depending on the experiment (**Figs. 3D, 5H**). The overhead camera served to track the centroid position of the fish in real time using the KNN background subtractor function from the openCV library. 60 seconds after the escape or disappearance, the virtual fish re-appeared on the screen and the process repeated for a total of three trials per fish. The number of escaping or disappearing virtual fish was randomized for experiments with differing numbers of escapees, with each fish experiencing a 1-, 3-, and 5-fish-escape event from the group of 5 total in the school (**Fig. 3F,G**). In the case of disappearing fish, they vanished at the onset of “escape”.

### Fish immobilization and holding chamber for *in vivo* two-photon calcium imaging

We modified the head-tethered preparation and holding chamber design from Zada et al. 2024 (ref ^18^). Briefly, adult *D. cerebrum* were transiently anesthetized with tricaine (120 mg/L MS-222 for 1-2 minutes) before removed from water and placed ventral side up on a 24x30x1 mm coverslip (12545B, Fisher Scientific). Two pieces of pre-cured SYLGARD wedges was fixed on the coverslip with UV-cured plastic (Bondic) to support the fish body from behind the eyes. 5-10% low-melting point agarose (UltraPureTM LMP Agarose, Invitrogen) was added to cover the body from swim bladder, and on the side of head in front of the SYLGARD pieces. Another horizontal strip UV-cured plastic was added to connect the coverslip and the agarose for more stability. Agarose from side of the eyes, gills, and the tip of the tail were removed with scalpel. Once all components were dried, fish was placed back into a holding fish and ventilated with oxygenated water to ensure gill movement. Those recovered and showed spontaneous breathing were then moved into the holding chamber under two-photon microscope.

The triangular chamber built from two monitors (HAMTYSAN, 7-inch 800x480 LCD). The two screens occupied ∼260 ° of visual space in azimuth (130 ° each side, with ∼10 ° gap in the front, and ∼90 ° gap behind the fish), and ∼105 ° of visual space in elevation (52.5 ° below and above). The chamber was filled with fish housing water (28°C, about 500 mL). The water level was maintained at about 1 cm above the midline of monitor height (10cm) by gravity flow of fresh fish housing water and a custom vacuum made from a bent borosilicate electrode (WPI) constantly removing excess water throughout the duration of the experiment.

The back wall and floors were made of transparent acrylic for IR lights and recording camera access. A 4.5 mm column of clear acrylic was used as a pedestal to position head-fixed fish at a standard positioning within this arena (intersection of each screen’s midpoint). Coverslips were suspended from the acrylic pedestal using two 3 x 1 mm magnets (Ethcool) that was embedded into the top of the pedestal and another one added when anchored.

### Visual stimuli and two-photon imaging

BonsaiRx was used to record the timing of microscope frame acquisitions, and display visual stimuli. Visual stimuli were displayed on the screens using cube mapping in BonVision^68^, where each screen displayed a viewport in a virtual environment. The virtual environment was composed of a 6000 mm^2^ plane with a smoothed white-noise pattern (“seafloor”). The fish’s position in this VR world was in the center (0,0 in x/y), 10 mm above the seafloor

For nonsocial stimuli, a dark sphere was programmed to appear in the front of fish in the virtual environment. After 1 sec, it either moves horizontally to the peripheral (sphere stimulus) or move toward the virtual camera (loom stimulus) over 5 sec (**Fig. 4B**). For the social stimuli, behavior clips of swimming and escaping *D. cerebrum* from both top-down and lateral view were processed to get realistic position changes of fish in 3-dimensional space (**Supplemental Fig. 6A**). Behavior tracking (121 and 200 Hz for the top-down and lateral side view) was resampled to the refreshing rate of monitors (60 Hz) and applied to spheres. The diameter and distance of the spheres in the virtual environment were calibrated to match their size with live fish. These two smaller dots swim from the back to front of the head-tethered fish in the VR environment over 6 sec (swim stimulus) or executed a downward and outward escape at the 3 sec mark and kept swimming farther (**Fig. 4B, Supplemental Fig. 6A**). All stimulus type are mainly rendered on one of the monitors, and a mirrored version is made to show the exact stimulus on the other side.

Each recording session started with a 1-minute baseline with only the seafloor was visible. We then used a trial structure to present each of the 6-sec stimulus type to head-tethered fish for 10 trials each in pseudorandomized order, with randomized 14-16 sec inter-trial intervals. All stimuli were displayed underneath the water surface of the recording arena and on the same side of monitor within the recording session (∼15 min).

Two-photon microscopy was performed with a ThorLabs Bergamo II multiphoton microscope controlled by ThorImageLS 4.3 and illuminated by an fs-pulsed 80-MHz Ti:S laser (MaiTai DeepSee; SpectraPhysics). GCaMP6s fluorescence was imaged with a 16x / 0.8W Nikon objective at an excitation wavelength of 930 nm and ≤10 mW excitation power. Frames of 1024 × 1024 pixels covered a field-of-view of 978x978 μm.

We imaged fish at a diagonal, to include as much of the brain as possible. Depth of the imaging plane was determined by maximizing the number of cells in optic tectum in the field of view. Two 2-plane and two single-plane imaging sessions were conducted for each fish if it survived the entirety experiments, where the stimuli came from each side of the visual field twice each. For two-plane imaging, we used fast Z mode in ThoImage4.3 to move across 50-90 μm at 5Hz. A transition cycle of 300 was pre-determined to make sure the Piezo has stabilized at the designated depths during the entirety of image acquisition of each plane. For the single-plane imaging, a 2-3 frame average was used to produce a final effective imaging rate at 5-7.6 Hz.

### Imaging analysis

We only proceeded with experiments where fish remained healthy at the end of recording and did not have substantial z-movements. For sessions with detected shifts in Z, we cropped the z-shifted frames if the remaining uncorrupted experimental length was more than half of the original timeseries.

Imaging data were motion corrected in Suite2p^58^, followed by cell extraction. Time series were inspected^70^ to ensure there was no z-motion contamination or slow drift, and only included neurons with continuous measurements and which were classified as a cell by Suite2p (“iscell”=1). A custom classifier was trained for more accurate cell extraction. The anatomical region each neuron located was further classified with manually defined boundaries. We excluded the first 5% of each imaging trial to avoid the influence of sound-evoked activity upon the initiation of scanning. Stimulus timing and calcium traces were aligned and resampled to a common 10 Hz sampling rate for further analysis. Then the fluorescence traces were detrended, smoothed with a 1-sec rolling mean, and z-scored. Z-scored fluorescence and the median x/y coordinates for each cell within these boundaries were used for further analysis.

We quantified stimulus responsive cells by calculating the response of a neuron by conducting t-test between the mean z-scored fluorescence during the 6-sec stimulus and the mean z-scored fluorescence in the 5-sec baseline pre stimulus. Neurons were identified as responsive to a certain stimulus type if the significance is *p*<0.05. All analyses were conducted using python code organized in Colab notebooks (see Code Availability).

For the support vector machine models, we only used experiments where there were at least 8 trials for all four stimulus types. The decoders were trained on the principal components (that explains 90% of variance) of neural activity of all responsive cells in each recording session. For longer sessions, 8 trials were randomly selected for each stimulus type and split with a stratified K-Fold cross-validator (8-fold). Twenty iterations of training and testing were performed.

### Statistics

Groups were tested for normality using the Shapiro-Wilk test. Non-parametric tests (Mann-Whitney U, Kruskal-Wallis H) were conducted if the Shapiro-Wilk *p* <0.05 for any group. Otherwise, parametric tests were used. *Post hoc* tests were conducted in main effect was *p*<0.05. All *post hoc* tests (Dunn’s test for nonparametric data, or t-tests for parametric data) were corrected for multiple comparisons with a Bonferroni correction. For nested data (neurons within imaging session within experimental groups), we determined the significance of pairwise comparisons using linear mixed-effects models, with imaging session identity (id) as a random variable. Exact tests and *p* values are reported in the figure legends.

### Data and code availability

Custom software (in Colab notebooks) and data will be made available upon publication.

